# Spatial heterogeneity of disease infection attributable to neighbor genotypic identity in barley cultivars

**DOI:** 10.1101/2025.04.22.650038

**Authors:** Iqra Akram, Lukas Rohr, Kentaro K. Shimizu, Rie Shimizu-Inatsugi, Yasuhiro Sato

## Abstract

Pest damage exhibits considerable spatial heterogeneity among individual plants in the field. While such spatial heterogeneity has often been treated as a nuisance in crop breeding trials, underlying biotic factors and loci remain poorly understood. To quantify the spatial variation in disease infection and associate it with neighboring genotypes, we applied two methods, Spatial Analysis of Field Trials with Splines (SpATS) and Neighbor Genome-Wide Association Study (Neighbor GWAS), to barley cultivars. Having compiled the CIMMYT Australia ICARDA Germplasm Evaluation (CAIGE) data, we first applied SpATS to three disease phenotypes such as the net form net blotch, spot form net blotch, and scald damage. This SpATS analysis showed extraneous phenotypic variation unexplained by smooth spatial trends, thereby leading us to focus on neighboring genotypes as an extraneous biological factor. We then applied the Neighbor GWAS model and found that neighbor genotypic identity explained 0.1–0.3 fractional variation in the three disease phenotypes. The Neighbor GWAS method also detected two significant or marginally significant variants on the barley 7H chromosome, which were associated with neighbor genotypic influence on the net form net blotch and scald damage. These variants were estimated to have beneficial effects that reduce disease damage by their allelic mixtures. Our findings suggest that neighbor genotypic identity can account for spatial variation in disease infection, providing a key to reduce pest damage by variety mixtures in field crops.

## Introduction

Natural and field-grown plants exhibit considerable spatial variation in their phenotypes, which are shaped by abiotic and biotic factors. Pest damage, such as pathogen infection (Rieux *et al*. 2014; Van Der Heyden *et al*. 2021) and insect herbivory (The Herbivory Variability Network *et al*. 2023), is especially heterogeneous among conspecific individual plants in the field. While spatial heterogeneity has been considered a nuisance in crop breeding trials (Rodríguez-Álvarez *et al*. 2018), extraneous spatial variation is sometimes shaped by biotic interactions. For example, neighboring genotypes are one of the biotic factors that can shape spatial heterogeneity of pest damage (Costa e Silva *et al*. 2017; Dahlin *et al*. 2018; Tamura *et al*. 2020; Pélissier *et al*. 2023). In close proximity, plant-plant interactions are driven by volatile communication among genotypes, allowing a variety mixture to be resistant to pests (Dahlin *et al*. 2018). Even when direct plant-plant communication is absent, resistant genotypes protect susceptible neighbors from pest migration (Tamura *et al*. 2020), or susceptible genotypes contrarily spread pests to resistant genotypes *vice versa* (Utsumi *et al*. 2011). Understanding the genetic architecture of conspecific neighbor effects may offer a way of pest control by variety mixtures (Mundt 2002; Dahlin *et al*. 2018; Montazeaud *et al*. 2022; Sato and Wuest 2024).

Barley (*Hordeum vulgare*) is an important cereal crop that is infected with various fungal pathogens worldwide. For example, *Pyrenophora teres* is a causal agent of net blotch symptoms in barley (Liu *et al*. 2011), in which *P. teres* f. *teres* and *P. teres* f. *maculata* induce different symptoms, namely, the net and spot form net blotch, respectively. In addition to *P. teres*, *Rhynchosporium secalis* is known as a causal agent of barley scald also known as ‘leaf blotch’ (Abang *et al*. 2006; Zhan *et al*. 2008). These fungi immediately develop spores on primary host plants under moderate climates and splash conidia by wind, leading secondary infection to another plant during the barley growing season (Zhan *et al*. 2008; Liu *et al*. 2011). To date, quantitative trait locus (QTL) mapping and genome-wide association studies (GWAS) have identified multiple loci associated with barley resistance to net blotch and scald damage (Tamang *et al*. 2015; Richards *et al*. 2017; Novakazi *et al*. 2019). Little is known, however, about the influence of neighboring genotypes and their associated variants.

To analyze the influence of neighboring genotypes on individual phenotypes, our previous study proposed a new method of genome-wide association study (GWAS), called “Neighbor GWAS” (Sato *et al*. 2021b). Inspired by the Ising model of magnetics, the Neighbor GWAS method expanded a standard mixed model to incorporate locus-wise interactions among neighboring genotypes. Owing to its locus-wise modeling, the Neighbor GWAS method can be applied to any randomized cultivation of multiple varieties, in which two alleles at a given locus are randomly distributed in space. As a randomized spatial arrangement has often been adopted in GWAS experiments (e.g., Cui *et al*. 2016; Sato *et al*. 2024; Behera *et al*. 2024), the Neighbor GWAS method is widely applicable to any plant GWASs. For instance, our previous study demonstrated the power of Neighbor GWAS to predict effective genotype mixtures that mitigated insect herbivory in *Arabidopsis thaliana*. This workflow can be applied to both individual- and plot-level data, as long as phenotypes are collected per genotype. Although the Neighbor GWAS method seems to suit agricultural GWAS data, this has not been applied to any crop.

In this study, we examined the influence of neighboring genotypes on disease infections, such as net blotch and scaled damage, in barley. Specifically, we aimed to address the following questions: (i) Was there extraneous spatial variation in disease infection? (ii) To what extent was the spatial variation explained by neighbor genotypic identity? (iii) Were there any significant variants associated with the neighbor genotypic effects on disease infection? To address these questions, we applied two methods, Spatial Analysis of Field Trials with Splines (SpATS) and Neighbor GWAS, to publicly available data collected by the CIMMYT Australia ICARDA Germplasm Evaluation (CAIGE) project (Trethowan *et al*. 2024).

## Materials & Methods

### Dataset

Barley phenotype and genotype data were obtained from the website of CAIGE (Trethowan *et al*. 2024) project via https://www.caigeproject.org.au/ (see also Table S1 for exact URLs). Specifically, DArT-derived SNP data for 807 lines were downloaded from the Gigwa database. Genotype data were imputed using BEAGLE version 5.1 (Browning *et al*. 2018) after excluding duplicated markers. Phenotype data were downloaded from the disease-screening webpage. Datasets were curated with the following three criteria: (1) spatial information (i.e., rows and ranges of field plots) was available; (2) phenotypes were recorded for multiple years; (3) more than 500 individuals were available for statistical analysis. As a result, we were able to retrieve three-year data on three phenotypes; spot form net blotch; net form net blotch; and scald damage recorded at Horsham, Australia (Fig. S1). End-point damage levels (score variables) for each year were used as the representative phenotypes for each disease. For each phenotype, the curated dataset included approximately 25,000 SNP markers and 500 individuals with 480 genotypes. The details of the sample size are presented in Table 1.

**Table 1.** Summary of curated data from the CAIGE barley disease trials.

| **Phenotype** | **#Plants<sup>†</sup>** | **#Genotypes** |  | **#SNPs** | **#SNPs** |
| --- | --- | --- | --- | --- | --- |
|  |  | **†** | **%Missing** | **(MAF=1%)** | **(MAF=5%)** |
| Net form net blotch | 508 | 481 | 32 | 20,578 | 14,568 |
| Spot form net blotch | 507 | 485 | 25 | 20,583 | 14,570 |
| Scald | 509 | 482 | 25 | 20,527 | 14,580 |

†Shown is the number after excluding individuals whose genotypes are unavailable (i.e., %Missing).

### Spatial Analysis of Field Trials with Splines (SpATS)

We used the Spatial Analysis of Field Trials with Splines (SpATS) (Rodríguez-Álvarez *et al*. 2018) to quantify the smoothing and extraneous spatial heterogeneity in the net form net blotch, spot form net blotch, and scald damage. The SpATS method employs a spline method to estimate spatial trends across two-dimensional space along rows and ranges according to their spatial proximity (Rodríguez-Álvarez *et al*. 2018). By modeling spatial trends, SpATS can also estimate the amount of phenotypic variation owing to genotypes and the spatial trends (Rodríguez-Álvarez *et al*. 2015), allowing us to quantify broad-sense heritability attributable to plants’ own genotypic effects. Extraneous spatial heterogeneity can also be visualized by calculating the row and column displacements for unexplained residuals (Gilmour *et al*. 1997) (see Figure 1 for example).

**Figure 1.**
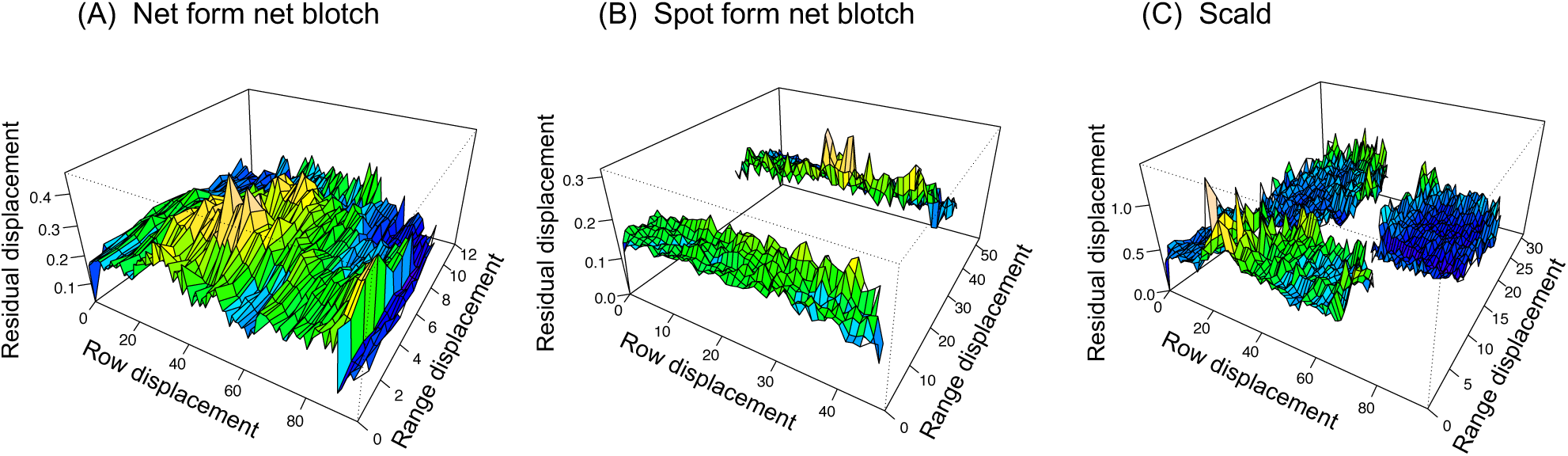
Sample variograms for visualization of spatial heterogeneity for net form net blotch (A), spot form net blotch (B), and scald (C). The X-axis represents the row displacement; the Y-axis represents the column displacement; and the Z-axis represents the residual displacement of disease severity between a pair of spatial positions (see Gilmour et al. 1997). The color gradient reflects disease severity levels where blue indicates low, green indicates moderate, and yellow indicates high severity.

To apply the SpATS method to barley data, we utilized the SpATS package (Rodríguez-Álvarez *et al*. 2018) implemented in R version 4.3.0 (R Core Team 2023), with the following arguments specified in the SpATS and other optional functions. Damage levels (score variable) for the net form net blotch, spot form net blotch, and scald were separately analyzed as a response variable. The study years were considered non-genetic covariates and were included as fixed effects. Genotypes (i.e., the name of cultivars) and spatial positions (i.e., row and range) were considered random effects; the former was specified as “genotype.as.random = TRUE” and the latter was specified as “random = row + range” within the SpATS function. The other parameters were set as default in SpATS and its auxiliary SAP function, which corresponded to a cubic B-spline and second penalty order (Rodríguez-Álvarez *et al*. 2018). The getHeritability function was also used to calculate a generalized broad-sense heritability *H*^2^(Oakey *et al*. 2006). The estimated spatial heterogeneity was visualized using the plot.variogram.SpATS function, which depicts sample variograms based on the displacements among rows, ranges, and residuals (Gilmour *et al*. 1997).

### Neighbor GWAS

#### Model description

We used the Neighbor GWAS method (Sato *et al*. 2021b) to quantify (i) the proportion of phenotypic variation explained (PVE) by neighbor genotypic effects and (ii) perform GWAS of neighbor genotypic effects on disease damage. In short, Neighbor GWAS consists of a two-factor linear mixed model that incorporates locus-wise identity (or similarity) of neighboring genotypes in addition to plants’ own genotypes as follows:

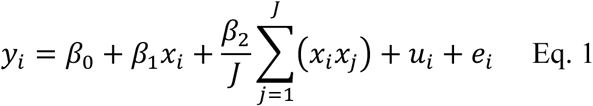

where *y*_*i*_ is a phenotype of *i*-th focal individual (or plot); β_0_ is an intercept; β_1_ is a fixed effect from plant’s own genotype at a focal locus *x*_*i*_; β_2_ is a fixed effect from genotypes of neighboring individuals (or plots) *x*_j_at the same locus (*j* up to the number of neighboring individuals *J*); *u*_*i*_ is a random effect; and *e*_*i*_ is a residual. For the two fixed effects β_1_ and β_2_, plant’s own genotypic values *x*_*i*_ are converted as −1, 0, and +1 for one homozygote, heterozygote, and the other homozygote, respectively. Accordingly, the mean allelic identity between the focal and neighboring plants ∑^*J*^_j=1_ (*x*_*i*_*x*_j_* /*J* takes from −1 (dissimilar) to +1 (similar), in which the sign of β_2_ corresponded to negative or positive effects of allelic identity on a focal plant’s phenotype *y*_*i*_ (Sato *et al*. 2021b). Therefore, statistical tests of β_2_ and its estimates enable GWAS with the inference for the positive or negative allelic interactions among neighboring plants. For the random effects *u*_*i*_, two variance-covariance matrices related to plant’s own and neighboring genotypes are considered as *u*_*i*_ ∼ Norm(0, σ_1_^2^*K*_1_ + σ_2_^2^*K*_2_). The two variance component parameters σ^2^ and σ^2^ respectively represents polygenic effects of plant’s own and neighboring genotypes on a phenotype *y*_*i*_, in which *K*_1_ is a kinship matrix and *K*_2_ represents neighbor genotypic similarity across a field (see page 8 of Sato *et al*. (2024) for the detailed definition of *K*_2_). The remaining phenotypic variation, i.e., residual, follows a normal distribution as *e*_*i*_ ∼ Norm(0, σ^2^I). Using Eq. 1, variation partitioning can be performed by estimating two variance components σ_1_^2^ and σ_2_^2^ (see “Phenotypic variation explained by neighboring genotypes” below) while GWAS can be performed by testing the fixed effect β_2_ (see “Genome-wide association study” below).

Standard GWAS was a subset of the Neighbor GWAS model (Eq. 1) when β_2_ and σ^2^ were set at 0. All the relevant methods of Neighbor GWAS are implemented as the rNeighbor GWAS package v1.2.4 (Sato *et al*. 2021b) in R, which depends on the gaston package (Perdry and Dandine-Roulland 2022) and uses its lmm.aireml and lmm.diago functions to solve mixed models and perform GWAS, respectively. Standard GWAS was also performed using Neighbor GWAS, which internally uses the gaston package of R (Perdry and Dandine-Roulland 2022). The theoretical details are described in Sato *et al*. (2021b).

#### Phenotypic variation explained by plants’ own and neighboring genotypes

We used the Neighbor GWAS package to estimate the variance components σ_1_^2^, σ_2_^2^, and σ_e_^2^ and examined how much spatial variation can be explained by plants’ own and neighboring genotypes. Net phenotypic variation explained (PVE) by neighboring genotypes can be quantified as PVE = [(σ_1_^2^ + σ_2_^2^)/(σ_e_^2^ + σ_2_^2^ + σ_1_^2^)] − *h*^2^, where *h*^2^ is a SNP heritability quantified by a standard GWAS model. To estimate the variance component parameters, we used 20k or 15k SNPs with the 1% or 5% cut-off threshold respectively applied to minor allele frequency (MAF) (see Table 1 for the exact number of SNPs). The latter cut-off percentage is often adopted in plant GWAS (Tibbs Cortes *et al*. 2021), while the former includes relatively rare but sometimes meaningful variants (Tibbs Cortes *et al*. 2021; Xu *et al*. 2023). The damage severity score (from 1 to 9) was separately analyzed for the net form net blotch, spot form net blotch, and scald damage. The endpoint damage score was used as the representative phenotype for each year. Non-genetic covariates were also considered for the differences among the three study years (2015, 2016, and 2017) and spatial positions (rows and ranges) of the field plots (see Fig. S1 for each year’s spatial arrangement). Along the rows and ranges, SpATS analysis detected the non-linear spatial trends of disease phenotypic values among field plots (see Results below); therefore, the first to fourth polynomials were considered for non-genetic covariates of rows and ranges to correct for the non-linear spatial patterns. Neighbor genotypic effects were assumed to be effective along rows/ranges and their diagonal positions (see Fig. 2A for a scheme). The maximum effective range was analyzed up to 3√2 Euclidean distance from focal plants (see the x-axes of Fig. 2B-D). This is because, when a spatial range is too broad, plant’s own genotypic value *x*_*i*_ and neighbor genotypic values ∑^*J*^ (*x*_*i*_*x*_j_*/*J* have severe co-linearity (Sato *et al*. 2021b, 2024).

**Figure 2.**
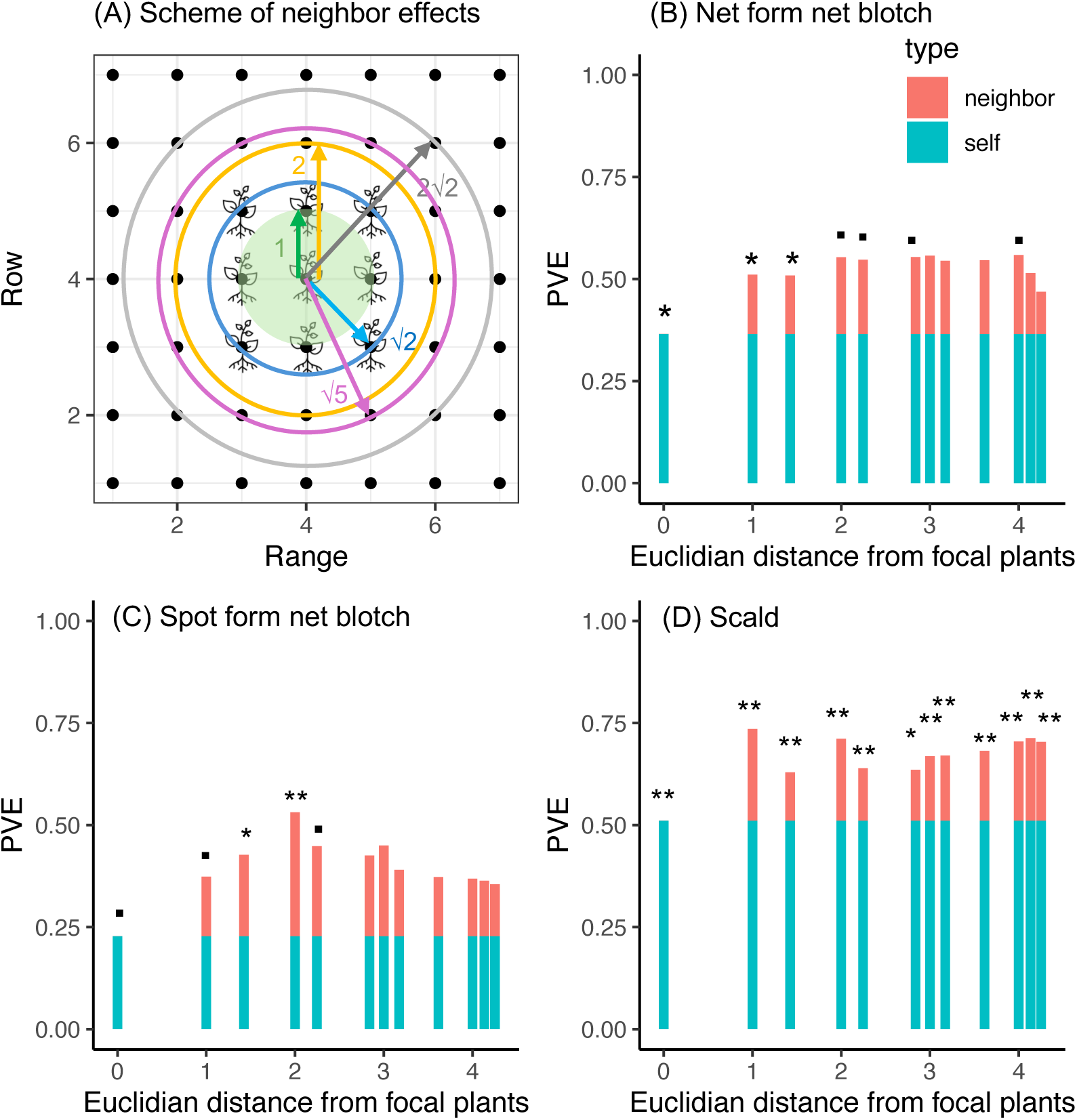
Analytical scheme and proportion of phenotypic variation explained (PVE) by plant’s own (self) and neighbor (neighbor) genotypic effects on the three phenotypes of disease infections. Panel (A) presents a schematic explanation of the range of neighbor genotypic effects assumed in the model, which corresponds to the distances on X-axes. Panels (B-D) display PVE for net form net blotch (B), spot form net blotch (C), and scald — (D). PVE at the distance of zero corresponds to a conventional SNP heritability (blue). Additional fractions explained by neighbor genotypic identity (red) indicate a net contribution of neighbor genotypic effects and are shown across the distances ≠ 0. Asterisks and dots indicate * *p* < 0.05; ** *p* < 0.01; ⋅ 0.05 < *p* < 0.1 with likelihood ratio test (see Table S3 for exact *p*-values).

To address this co-linearity, likelihood ratio tests were performed for Eq. 1 from simpler to complex models following Sato *et al*. (2021b). The actual spatial arrangement and number of neighbors are summarized in supplementary materials (Fig. S1 and Table S2).

#### Genome-wide association study (GWAS)

To narrow down genomic regions associated with neighbor genotypic effects on disease infection, we performed GWAS of the neighbor effect coefficient β_2_ and depicted Manhattan plots. The target phenotype and non-genetic covariates were the same as those used in the PVE analysis. Each SNP was tested after diagonalization on a weighted kinship matrix *K*′ = σB^2^*K*_1_ + σB^2^*K*_2_ (see Sato *et al*. 2021b for details). Same as the PVE analysis above, GWAS was repeated up to 3√2 Euclidean distance from focal plants and separately performed for the three phenotypes, i.e., the net form net blotch, spot form net blotch, and scald damage. Standard GWAS was also performed to test whether genomic regions overlapped between plant’s own and neighbor genotypic effects. We determined the statistical significance at *p* = 0.05 and the marginal significance between *p* = 0.05 and 0.1 throughout the present study. The inflation of *p*-values was diagnosed using quantile-quantile (QQ) plots. To check whether any genes were located near our focal SNPs, we referred to the genome annotation of Barley Morex V3 (Mascher *et al*. 2021) using the GrainGenes database (https://wheat.pw.usda.gov/GG3/genome_browser).

## Results

### Spatial heterogeneity of disease infection

We used SpATS to distinguish genetic and spatial variation in the three disease phenotypes — namely, net form net blotch, spot form net blotch, and scald damage — among barley cultivars. The generalized broad-sense heritability *H*^2^ (Oakey *et al*. 2006) was estimated as 0.78 for the net form net blotch; 0.68 for spot form net blotch; and 0.65 for scald damage, indicating that the moderate to high variation could be explained by plants’ own genotypes. Besides these genetic variations, SpATS analysis quantified spatial variation along the rows and columns of the field plots (Fig. S1; Table 2). The net form net blotch exhibited considerable spatial variation along the row position, while the spot form net blotch showed less spatial variation (Table 2). The scald damage exhibited substantial spatial variation along both the rows and ranges (Table 2). Nevertheless, all of these diseases had substantial residuals unexplained by genotypes and smooth spatial trends (Table 2). Such extraneous variation was displayed for unexplained residuals along the rows and columns, such that sample variograms presented a wavy pattern for each disease (Fig. 1A-C). These results showed that the substantial spatial heterogeneity of disease infection was unexplained by plants’ own genotypes and smooth spatial trends, leaving room to incorporate other factors, such as neighboring genotypes.

**Table 2.**
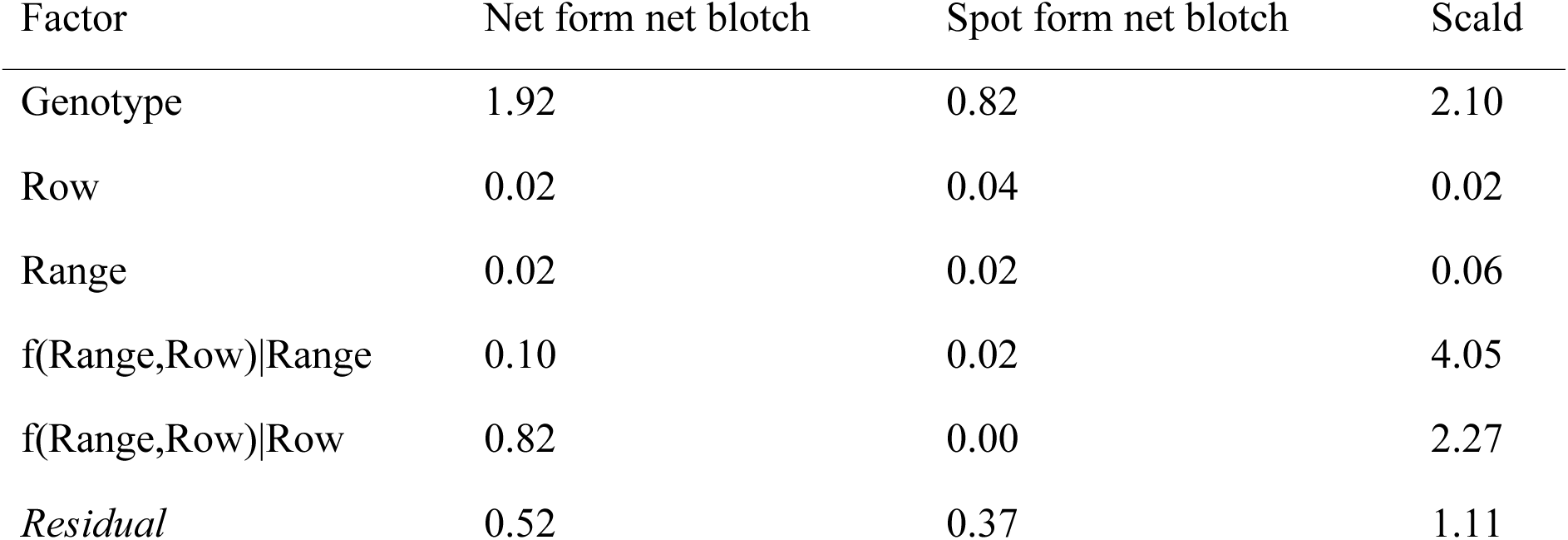
SpATS results showing variances for plant genotypes and 2D-spatial positions (range and rows). The estimated variance is shown for each random factor and phenotype. f(Range,Row)|Range or Row indicates a smoothing factor for the range and row positions.

### Phenotypic variation explained by plants’ own and neighboring genotypes

Subsequently, we asked to what extent the spatial heterogeneity could be explained by neighbor genotypic effects (Fig. 2A; Table S3). To address this question, we calculated phenotypic variation explained by neighbor genotypic identity using the Neighbor GWAS model with the rows and ranges incorporated as non-genetic covariates. All the three phenotypes of disease damage exhibited >0.2 SNP heritability *h*^2^ regarding plant’s own genotypic effects (blue bars in Fig. 2B-D). The net form net blotch showed 0.37 heritability at the 5% significance level (PVE at zero distance equivalent to *h*_SNP_^2^ = 0.366, *X*^2^ = 5.01, *p* < 0.05: see Table S3 for exact *p*-values). The spot form net blotch showed 0.23 heritability but at the marginally significance level (*h*_SNP_^2^ = 0.228, *X*^2^ = 2.91, *p* < 0.1). The scald damage showed 0.59 heritability at the 5% significance level (*h*_SNP_^2^ = 0.591, *X*^2^ = 14.3, *p* < 0.001).

These SNP-based narrow-sense heritabilities were lower than the broad-sense heritability described above, but this was also the case for previous studies on barley diseases (Zerihun *et al*. 2019; Kunze *et al*. 2024). These results confirmed that our SNP-based analysis was able to detect heritable variation in disease damage.

More remarkably, we detected significant contributions of neighboring genotypes to all the three diseases (red bars in Fig. 2B-D; likelihood ratio test *p* < 0.05 see Table S3 for exact test statistics and *p*-values). Specifically, the influence of neighboring genotypes on the net form net blotch was significant (*p* < 0.05) up to the second nearest neighbors and remained marginally significant (*p* < 0.1 and *p* > 0.05) at some scales up to the fourth nearest neighbors (Fig. 3B and Table S3A). The spot form net blotch was significantly influenced by the first and second nearest neighbors, though this influence became non-significant for distant neighbors (Fig. 2C and Table S3B). The scald damage showed the most significant patterns among the three disease phenotypes, such that the influence of neighboring genotypes remained significant across a space (Fig. 2D and Table S3C). These significant contributions of neighboring genotypes to the three disease phenotypes explained 0.1 to 0.3 faction of the total phenotypic variation (red bars in Fig. 2). Similar patterns were found even when the cut-off value of MAF was changed to 5% (Table S4A-C). These PVE analyses showed that a significant fraction of damage variation was attributable to neighbor genotypic identity, which led us to further ask whether major-effect SNPs accounted for this damage variation.

**Figure 3.**
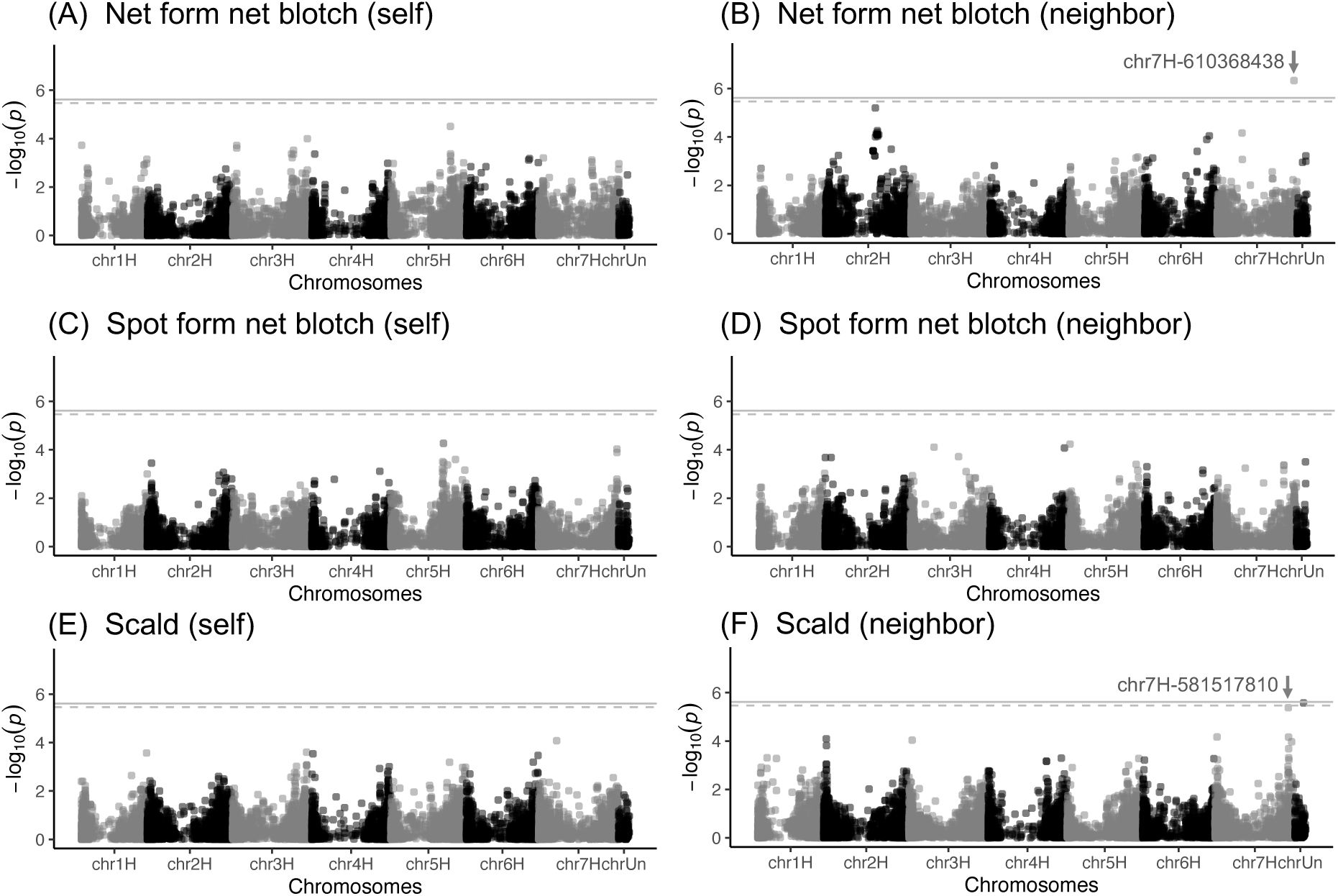
GWAS Manhattan plots for plant’s own or neighbor genotypic effects on the three phenotypes of disease infections in barley cultivars. Left (A, C, and E) and right (B, D, and F) panels display plant’s own (self) or neighbor (neighbor) genotypic effects, respectively. Target phenotypes are the net form net blotch (A-B), spot form net blotch (C-D), and scald (E-F). SNPs with MAF >1% are plotted against chromosomal positions, with solid and dashed horizontal lines indicating genome-wide Bonferroni threshold at *p* = 0.05 with MAF cut-off values at 1% and 5%, respectively. “chrUn” means ‘chromosome unassigned’ in the reference genome. Results at 4 Euclidean distances are shown for neighbor genotypic effects (B, D, and F).

### Genome-wide association study

Lastly, we used Neighbor GWAS to detect loci associated with neighbor genotypic effects on the three phenotypes of disease infection at spatial scales up to the third diagonal neighbors (Euclidean distance ≤ 3√2) (Fig. 3B, D and F; Fig. S2D-F). We detected two significant SNPs associated with neighbor genotypic effects at the spatial scale of the fourth nearest neighbors (i.e., Euclidean distance = 4 from focal plants). One of these two significant SNPs was detected for the net form net blotch, and the other was for scald damage (Fig. 3B and F; Fig. S2D and F). The significant SNP of neighbor genotypic effects on the net form net blotch was located on the tip of the 7H chromosome (Chr7H at 610368438: Fig. 3B; Fig. 4A). This significant SNP exhibited a positive sign of neighbor effect coefficient (β_2_= 2.77, MAF = 0.109, raw *p*-value = 4.66e-07, Bonferroni-corrected *p*-value = 0.0096 when MAF cut-off at 1%), indicating that this SNP had beneficial allelic interactions to prevent the disease spread among neighboring genotypes. The other significant SNP, which was detected for the scald damage, was assigned as an unanchored chromosome (ChrUn at 68152334: β_2_= 3.261841, MAF = 0.085, raw *p*-value = 2.67e-06, Bonferroni-corrected *p*-value = 0.034 when MAF cut-off at 5%: Fig. 3F); therefore, its QTL position was unknown. In addition to the two significant SNPs, a marginally significant SNP for the scald damage was found on the 7H chromosome (Chr7H at 581517810: Fig. 3F; Fig. 4B). This SNP had a positive sign of the neighbor effect coefficient (β_2_= 3.24, MAF = 0.098, raw *p*-value = 4.19e-06, Bonferroni-corrected *p*-value = 0.086 when MAF cut-off at 1%), indicating beneficial allelic interactions at this locus to prevent the disease spread among neighboring genotypes. These findings suggest the potential existence of QTLs responsible for beneficial genotype-genotype interactions that mitigate disease spread through long-range neighbor effects.

**Figure 4.**
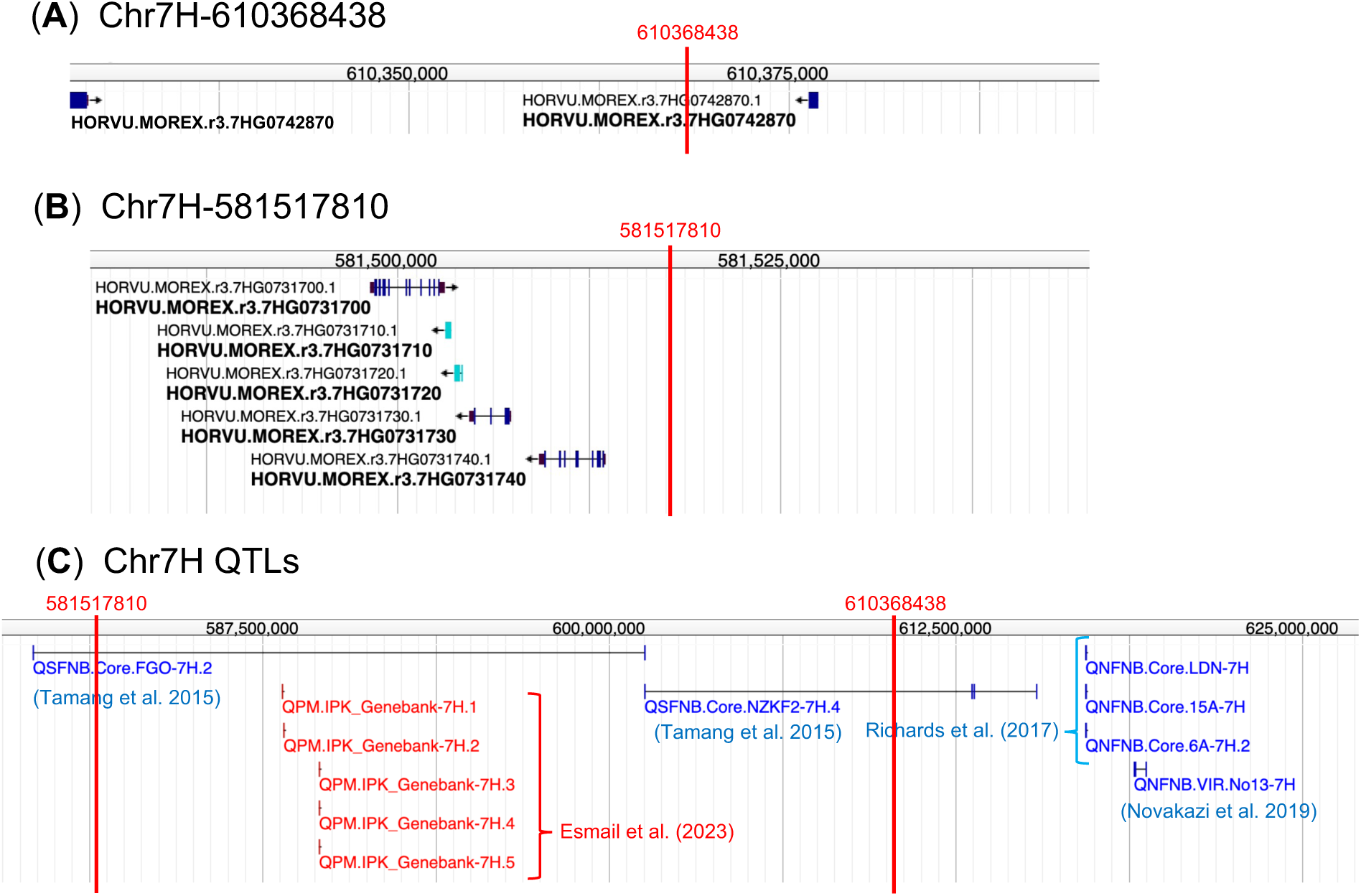
Candidate genes and quantitative trait loci (QTLs) near focal SNPs on the barley 7H chromosome. (A) The nearest two genes to a significant SNP associated with neighbor genotypic effects on the net form net blotch shown (cf. Fig. 3B). (B) The five nearest genes to a marginally significant SNP associated with neighbor genotypic effects on the scald damage (cf. Fig. 3F). (C) QTLs located near 581517810 and 610368438 bp genomic positions on the 7H chromosome. Numbers above the horizontal axes indicate base pairs. Genome annotations were obtained from the GrainGenes database (https://wheat.pw.usda.gov/GG3/genome_browser).

Additionally, we conducted a standard GWAS to examine the overlap of QTLs between plants’ own and neighbor genotypic effects (Fig. 3A, C and E; Fig. S2A-C). While this standard GWAS did not find any significant SNPs for the three diseases (*p* > 0.1 after Bonferroni correction: Fig. 3A, C and E), a weak peak was found on the tip of the 7H chromosome for the spot form net blotch (SNPs on Chr7H at 621031815, raw *p*-value = 1.297e-04; and at 621053989, raw *p*-value = 9.326e-05: Fig. 3C). Within 25 Mb of these SNPs, we found known QTLs responsible for barley resistance to the spot form net blotch (Tamang *et al*. 2015) and net form net blotch (Richards *et al*. 2017; Novakazi *et al*. 2019) (Fig. 4C). Together with previous evidence, the results of standard GWAS suggest that the same gene function can mediate both plants’ own and neighbor genotypic effects on disease infection (see also Discussion “Genetic architecture of neighbor effects” below).

## Discussion

Our analysis revealed that substantial fractions of spatial heterogeneity were attributable to neighboring genotypes in barley disease. Similar studies have reported the influence of neighboring genotypes on the other individuals’ phenotypes in *Arabidopsis thaliana* (Montazeaud *et al*. 2023; Sato *et al*. 2024), *Eucalyptus globulus* (Costa e Silva *et al*. 2013; Costa e Silva and Kerr 2013; Costa e Silva *et al*. 2017), and durum wheat *Triticum turgidum* ssp. *durum* (Montazeaud *et al*. 2022). While these previous studies also quantified the phenotypic variation attributable to neighboring genotypes, few have identified genetic variants associated with these effects (Montazeaud *et al*. 2022). In this context, our Neighbor GWAS analysis detected a significant SNP associated with neighbor genotypic effects on the net blotch damage. This is the first report for Neighbor GWAS to detect a significant QTL and, to our knowledge, one of few reports on significant QTLs associated with neighbor genotypic effects on agriculturally important traits.

### Genetic architecture of neighbor effects

Through marker-trait associations, GWAS can provide a way to screen candidate genes near significant SNPs (Korte and Farlow 2013; Tsuchimatsu *et al*. 2020; Tibbs Cortes *et al*. 2021). On the tip of the 7H chromosome, our analysis found a significant SNP at 610368438 for the net form net blotch and a marginally significant SNP at 581517810 for scald damage (Fig. 3B and F). Within 50 kb of the 610368438 position, we found the two loci, HORVU.MOREX.r3.7HG0742860 and HORVU.MOREX.r3.7HG0742870 (Fig. 4A), which encode protein kinases. The other SNP at 581517810 was located within 25 kb near the five loci (Fig. 4B), including HORVU.MOREX.r3.7HG0731740, HORVU.MOREX.r3.7HG0731730, and HORVU.MOREX.r3.7HG0731700 which encode a SCO1-like protein, lipid transfer protein, and transducin/WD40 repeat protein, respectively. Among them, protein kinases and WD40 repeat proteins are known to be involved in various mechanisms of abiotic and biotic resistance in barley and wheat (Ruiz-Roldán *et al*. 2001; Kong *et al*. 2015; Liu *et al*. 2019; Yan *et al*. 2023). These are candidate genes for future studies after GWAS, such as functional analyses using single-gene mutants for mechanistic understanding (e.g., Tsuchimatsu *et al*. 2020).

Evidence of disease resistance loci is accumulating for important crops such as barley and wheat (Jung *et al*. in press; Seeholzer *et al*. 2010; Keller *et al*. 2018; Walkowiak *et al*. 2020; Shimizu *et al*. 2021; Sotiropoulos *et al*. 2022). Besides candidate genes, previous studies on barley cultivars reported several disease-associated QTLs for the tip of the 7H chromosome (Tamang *et al*. 2015; Richards *et al*. 2017; Novakazi *et al*. 2019; Clare *et al*. 2020; Esmail *et al*. 2023) near the 581517810 and 610368438 bp positions (Fig. 4C). While our GWAS peaks were relatively weak regarding plant’s own genotypic effects, overlapping QTLs suggest that the same gene function can mediate both plants’ own and neighbor genotypic effects on disease resistance. For example, Tamang *et al*. (2015) performed GWAS using 2,062 barley accessions and identified a novel QTL associated with the damage caused by the New Zealand strain NZKF2 of the spot form net blotch. This QTL region ranged from 601275881 to 615403763 on the 7H chromosome (Tamang *et al*. 2015), including the SNP position of 610368438, for which we found long-range neighbor genotypic effects on the net form net blotch (Fig. 4C). This QTL region also encompassed five SNP markers associated with powdery mildew resistance as reported by Esmail *et al*. (2023) (see Fig. 4C). The other QTL reported by Tamang *et al*. (2015) ranged from 579216453 to 601275986 on 7H chromosome (Fig. 4C), encompassing the marginally significant SNP at 581517810, for which we found long-range neighbor genotypic effects on the scald damage. These QTLs may be a clue to design variety mixtures to prevent disease spread by optimizing genotype-genotype interactions.

The influence of neighboring genotypes on the disease infection of another plant could occur through disease dispersal from one plot to another. All the three fungal agents *P. teres* f. *teres*, *P. teres* f. *maculata*, and *R. secalis*, splash their conidia and accordingly cause a secondary infection of net form net blotch, spot form net blotch, and scald damage, respectively. We found that the magnitude of the neighbor genotypic effects differed among the three diseases, which might be attributed to two possibilities. One possibility is the spatial arrangement, because the least significant phenotype, spot form net blotch, has a spatial gap among years (Fig. 1B; Fig. S1B, E, and H). This might increase data heterogeneity and thus made the statistical power low for the spot form net blotch. The other possibility concerns the biological aspect of the fungal life cycle and dispersal modes. While few studies compared infection processes between *P. teres* f. *teres* and f. *maculata* in detail, Van Den Berg and Rossnagel (1990) reported a shorter infection time by *P. teres f. maculata* than *P. teres f. teres* on barley. Such differences in the fungal developmental rate might differentiate the influence of neighboring genotypes on disease infection.

### Applicability and limitation

In order to infer neighboring genotype-genotype interactions, our analysis showed the effective use of open data collected from a randomized block design of many plant genotypes. As the randomized block design is often employed to conduct GWAS (Cui *et al*. 2016; Sato *et al*. 2024; Behera *et al*. 2024), there should be other available data on various plant species. In this context, Neighbor GWAS does not require manipulative experiments and thus widens the opportunity to study the genetic architecture of genotype-genotype interactions using open data (Sato and Wuest 2024). Meanwhile, it should be noted that open data may not always be complete. For instance, the present barley data exhibited a rectangular arrangement with irregular intervals along rows and ranges in the field (Fig. S1). Detailed metadata, such as the physical distance between individual plots or plants, were not found in the open barley data. The shortage of basic information could have made the interpretation of Neighbor GWAS difficult. To solve this issue, pattern-based analyses, such as the P-spline method in SpATS (Rodríguez-Álvarez *et al*. 2018), were used as complementary tools to distinguish between smooth and extraneous spatial variation. A joint use of pattern- and process-based modeling would be effective in overcoming potential limitations regarding data availability.

To separate neighbor genotypic effects from non-genetic spatial trends, we incorporated polynomials of the rows and range positions as covariates into Neighbor GWAS. Despite the inclusion of these additional covariates, we detected an influence of neighboring genotypes on barley diseases. This strategy may be applicable to other GWAS data where a number of crop genotypes are cultivated at an individual or plot level across a large field. However, the number of orders that should be considered for row and range polynomials remains unknown. When ruling out spatial heterogeneity as a nuisance, neighbor genotypic effects are confounded and are thus difficult to separate from spatial trends (Sato *et al*. 2021a).

Integration of the two mixed models, SpATS and Neighbor GWAS, is challenging for GWAS implementation but is needed to fully distinguish spatial trends and neighbor genotypic effects on a phenotype.

## Conclusion

By harnessing open data, we detected significant phenotypic variation and genetic variants associated with neighbor genotypic effects on disease infection in barley. Such neighbor genotypic effects are linked to the effects of genotype mixtures on pest damage (Sato *et al*. 2024), providing a promising way for integrated pest management by mixed planting (Mundt 2002; Tooker and Frank 2012). Unlike in a previous study on polygenic traits (Sato *et al*. 2024), the identification of significant QTLs may enable us to optimize population-level pest damage with a few loci being targeted (Wuest *et al*. 2021; Sato and Wuest 2024). For barley cultivars, we detected a significant SNP that had beneficial effects to reduce disease damage by allelic mixtures, indicating the potential prevention of disease spread by allelic mixtures. Beyond genotype mixture, allelic mixture is particularly suitable for crop varieties because intraspecific varieties can be subjected to breeding by crossing each other (Wuest *et al*. 2021; Sato and Wuest 2024). Further studies are needed to validate these effects by comparing allelic monoculture and mixture at the candidate locus.

## Data availability

All source codes and input data are available at GitHub (https://github.com/yassato/caige_barley). The full list of SNP positions and -log_10_(*p*) association scores is also available in the GitHub repository as a binary format of R language. Supplementary Figure S1 and S2 present field plot positions and GWAS QQ plots, respectively. Supplementary Table S1 shows the exact URLs for the original input data. Supplementary Table S2 shows the actual number of neighboring plants. Supplementary Tables S3 and S4 include the exact *p*-values for the likelihood ratio tests for PVE.

## Acknowledgements

The authors appreciate all efforts made by the data collectors of the CAIGE project. We especially thank Richard Trethowan and Amit Singh for their responses to our inquiry. The computing resource was provided by Human Genome Center at the University of Tokyo (http://sc.hgc.jp/shirokane.html).

## Supplementary Materials

**Figure S1.**
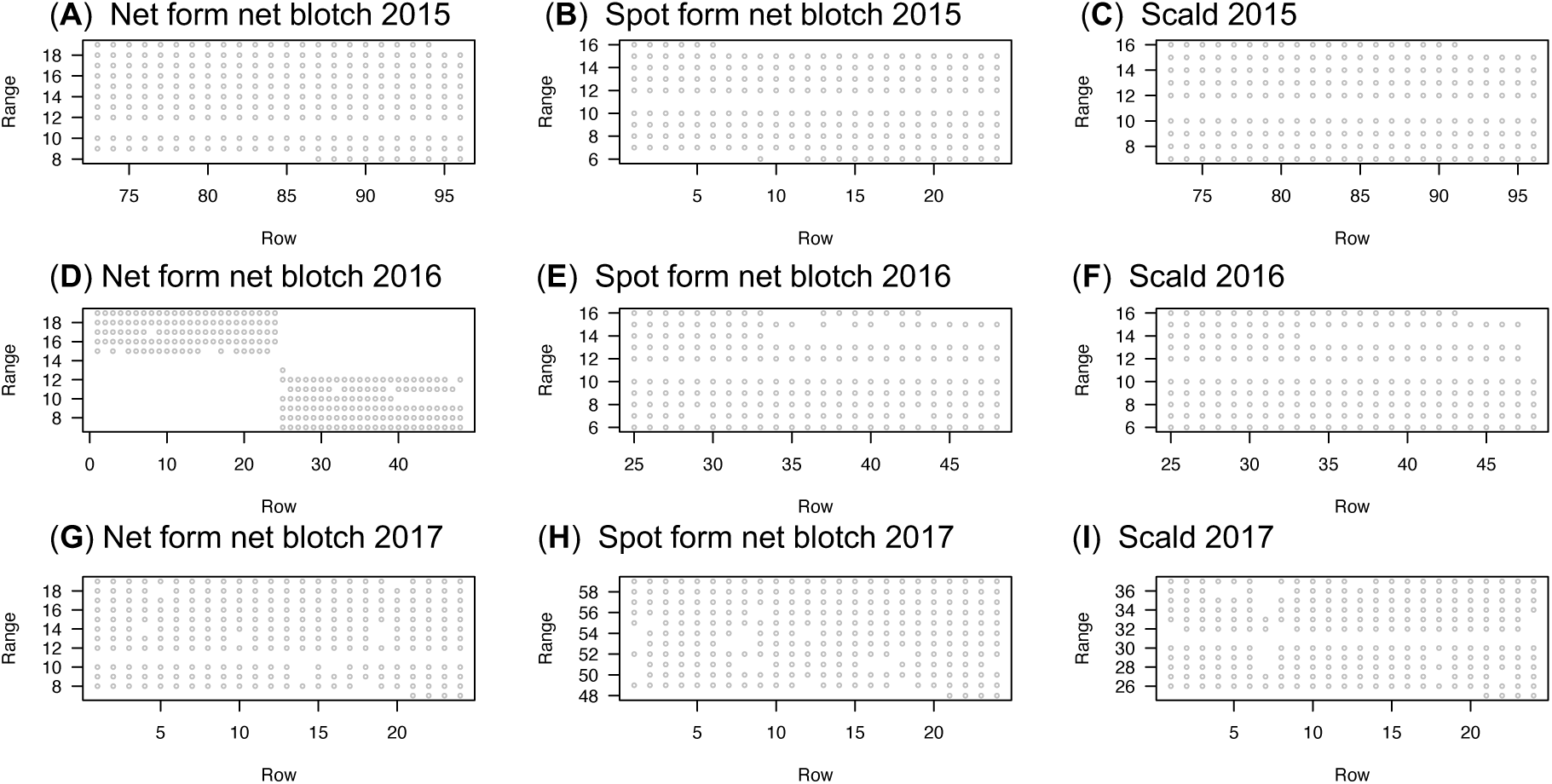
Field plot positions along the range and rows for each year and phenotype. A single circle corresponds to a plot. Neighbor genotypic similarity was calculated within a year — namely, we did not assume neighbor interactions among years. Yearly differences were included as non-genetic covariates for each phenotype. See also “Materials & Methods” for details of statistical analyses.

**Figure S2.**
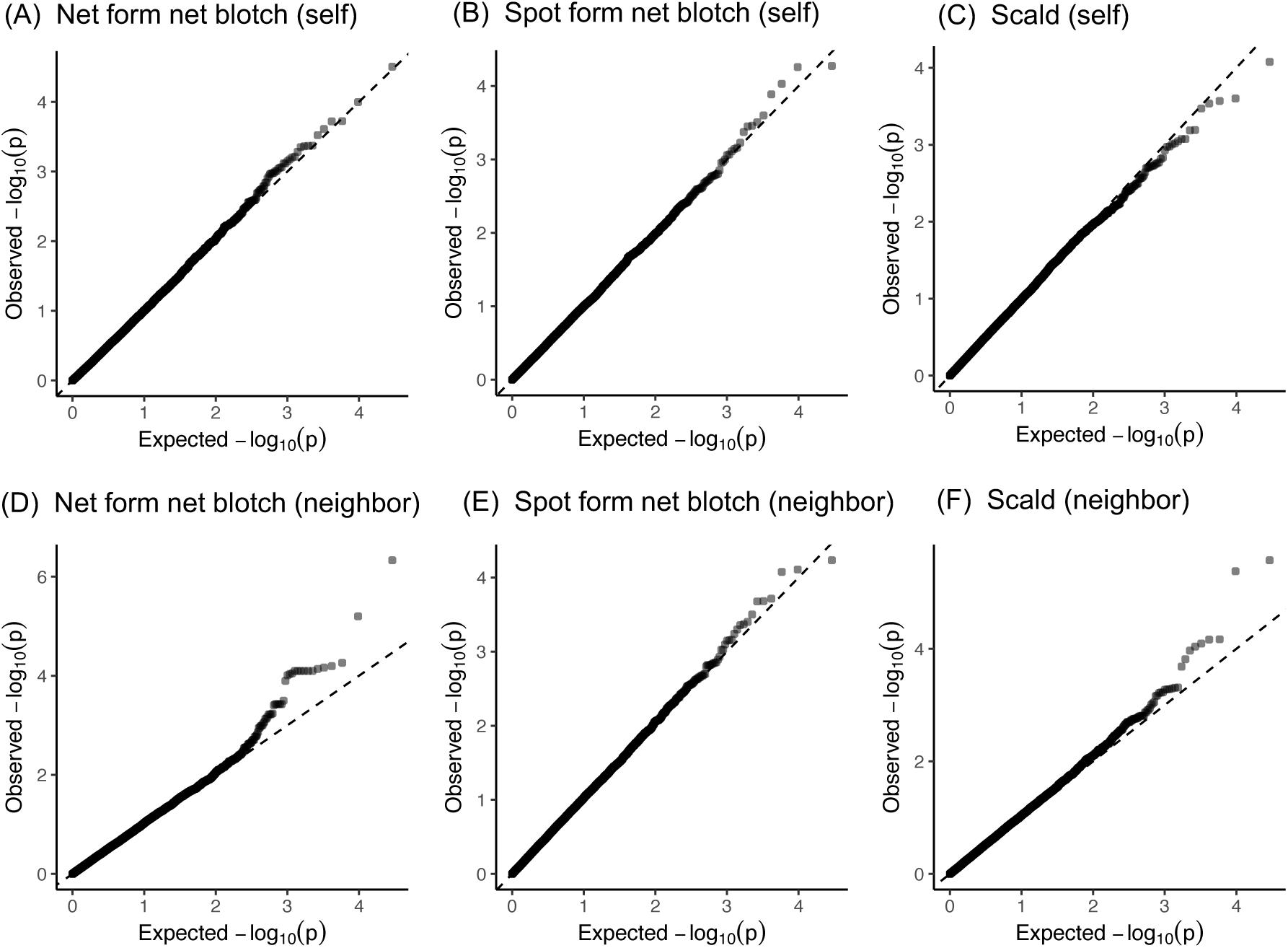
Quantile-Quantile (QQ) plots of plant’s own or neighbor genotypic effects on the three phenotypes of disease infections in barley cultivars. The upper (A, B, and C) and lower (D, E, and F) panels display plant’s own (self) or neighbor (neighbor) genotypic effects, respectively. Target phenotypes are the net form net blotch (A-D), spot form net blotch (C-E), and scald (C-F). The observed -log_10_(*p*) association scores are plotted against expected values. The dashed lines indicate random expectation as *y* = *x*.

**Table S1.**
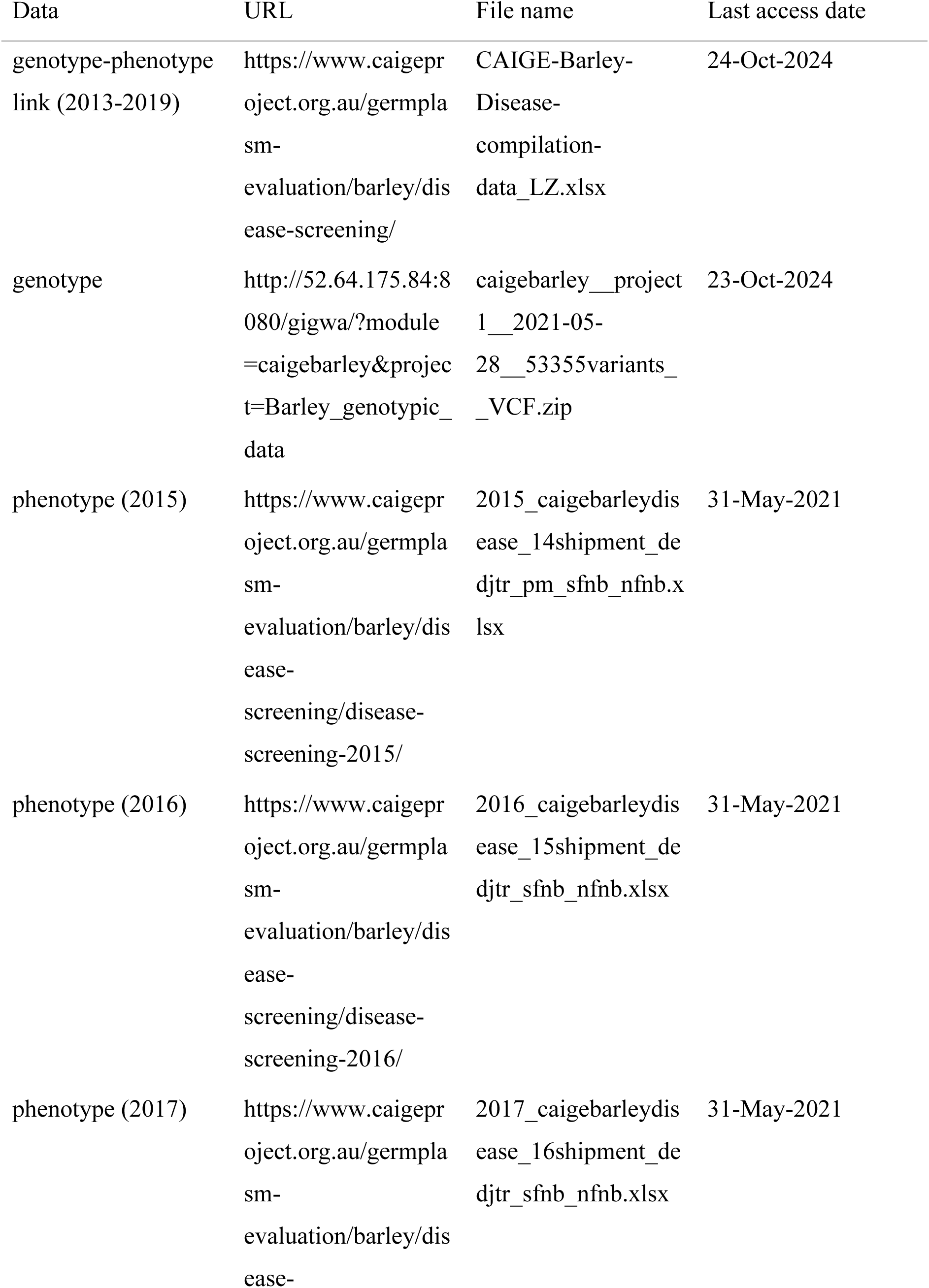
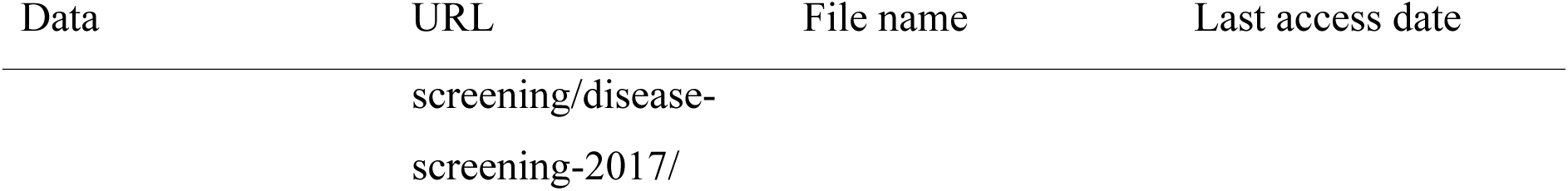
Exact links to the study data within the CAIGE website.

**Table S2.** The theoretical and actual number of neighboring plants for the net form net blotch, spot form net blotch and scald damage. The average number of neighbors is shown for each Euclidean distance from focal plants. See also Figure S1 for actual data points.

| **Distance** | **theoretical max.** | **Net form net blotch** | **Spot form net blotch** | **Scald** |
| --- | --- | --- | --- | --- |
| 0.000 | 0 | 0 | 0 | 0 |
| 1.000 | 4 | 2.8 | 3.1 | 3 |
| 1.424 | 8 | 4.9 | 5.6 | 5.3 |
| 2.000 | 12 | 7.3 | 8.3 | 8 |
| 2.246 | 20 | 11 | 12.8 | 12.3 |
| 2.838 | 24 | 12.7 | 14.8 | 14.2 |
| 3.000 | 28 | 14.9 | 17.3 | 16.6 |
| 3.172 | 36 | 18.4 | 21.4 | 20.5 |
| 3.616 | 44 | 21.5 | 25.1 | 23.9 |
| 4.000 | 48 | 23.8 | 27.2 | 26 |
| 4.133 | 56 | 27.4 | 30.8 | 29.4 |
| 4.253 | 60 | 28.8 | 32.5 | 31 |

**Table S3.**
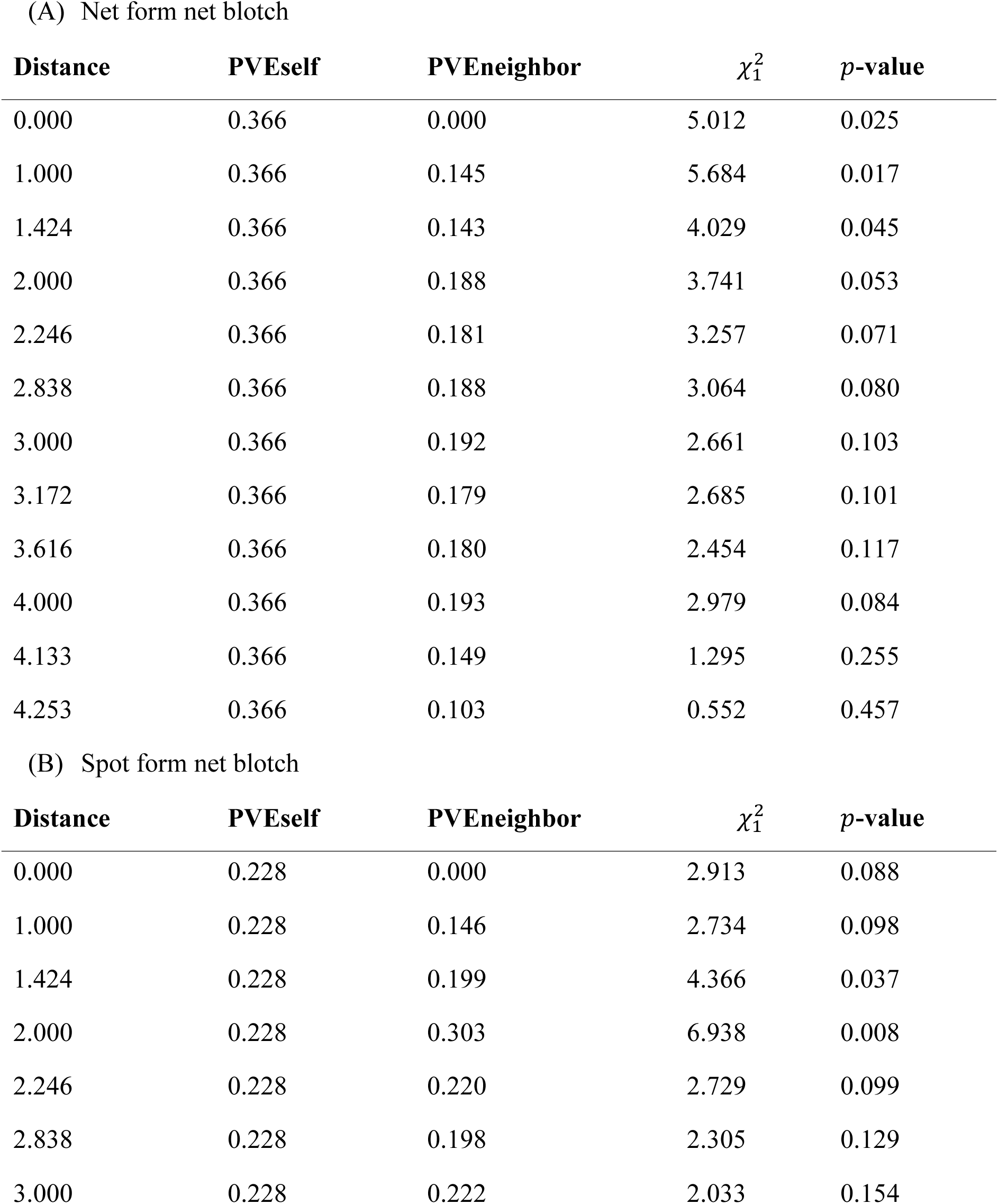
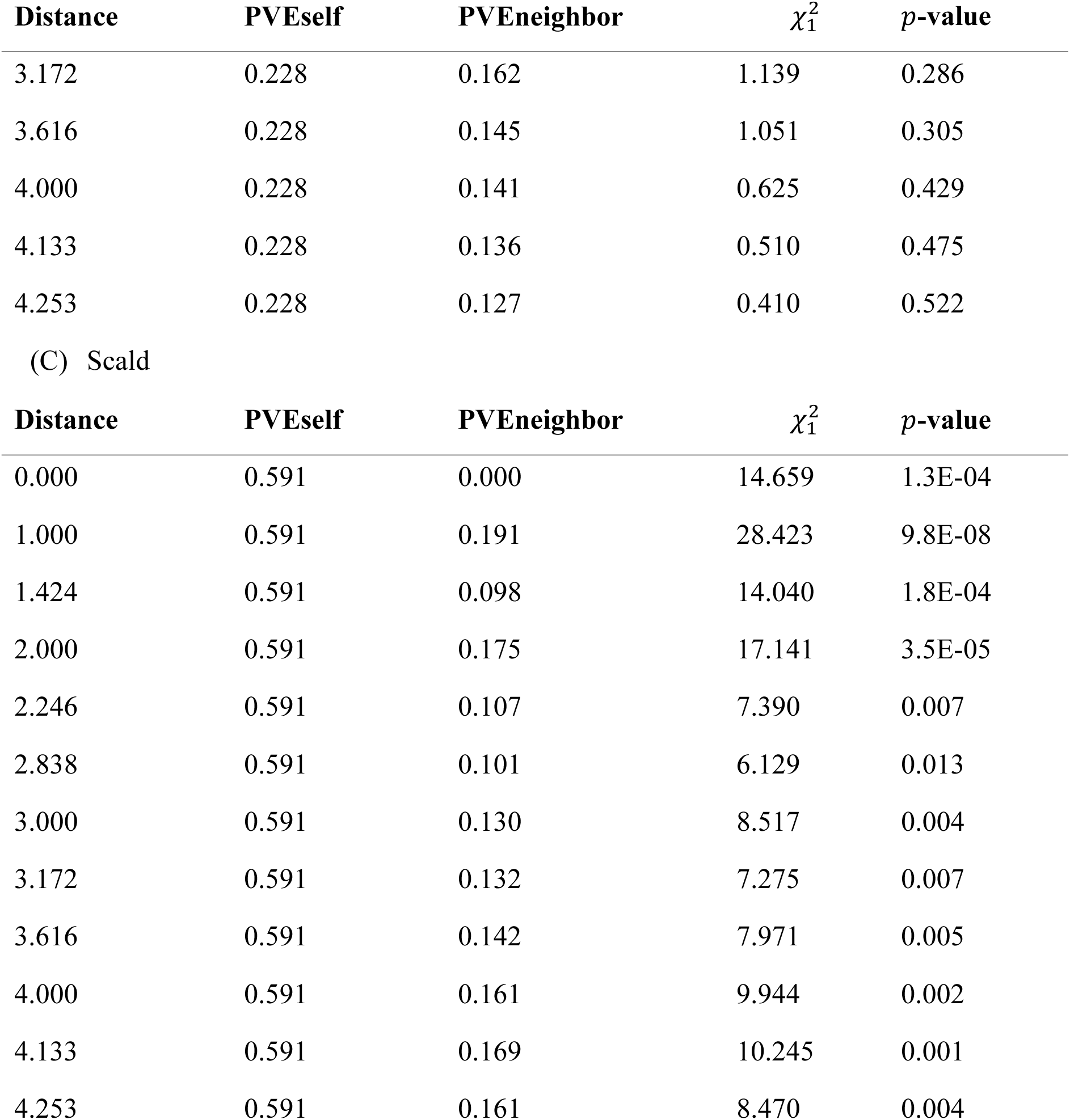
The proportion of phenotypic variation explained (PVE) by plant’s own (PVEself) or neighbor (PVEneighbor) genotypes with MAF cut-off at 1%. For distance ≠ 0, likelihood ratio tests based on *X*^2^-statistic were used to determine *p*-values of Neighbor GWAS models over a standard GWAS model. For distance = 0, likelihood ratio tests were performed over a null model, which made PVEself equivalent to a SNP heritability (see also Materials & Methods “Phenotypic variation explained by neighboring genotypes”).

**Table S4.** The proportion of phenotypic variation explained (PVE) by plant’s own (PVEself) or neighbor (PVEneighbor) genotypes with MAF cut-off at 5%. For distance ≠ 0, likelihood ratio tests based on *X*^2^-statistic were used to determine *p*-values of Neighbor GWAS models over a standard GWAS model. For distance = 0, likelihood ratio tests were performed over a null model, which made PVEself equivalent to a SNP heritability (see also Materials & Methods “Phenotypic variation explained by neighboring genotypes”).

| Distance | PVEself | PVEneighbor | $\chi^2_1$ | $p$-value |
| --- | --- | --- | --- | --- |
| 0.000 | 0.296 | 0.000 | 4.878 | 0.027 |
| 1.000 | 0.296 | 0.300 | 5.782 | 0.016 |
| 1.424 | 0.296 | 0.310 | 3.977 | 0.046 |
| 2.000 | 0.296 | 0.433 | 3.532 | 0.060 |
| 2.246 | 0.296 | 0.457 | 3.019 | 0.082 |
| 2.838 | 0.296 | 0.483 | 2.942 | 0.086 |
| 3.000 | 0.296 | 0.513 | 2.378 | 0.123 |
| 3.172 | 0.296 | 0.497 | 2.389 | 0.122 |
| 3.616 | 0.296 | 0.501 | 2.340 | 0.126 |
| 4.000 | 0.296 | 0.513 | 2.793 | 0.095 |
| 4.133 | 0.296 | 0.424 | 1.091 | 0.296 |
| 4.253 | 0.296 | 0.268 | 0.399 | 0.528 |

(B) Spot form net blotch
| Distance | PVEself | PVEneighbor | $\chi^2_1$ | $p$-value |
| --- | --- | --- | --- | --- |
| 0.000 | 0.166 | 0.000 | 2.598 | 0.107 |
| 1.000 | 0.166 | 0.145 | 2.449 | 0.118 |
| 1.424 | 0.166 | 0.207 | 4.117 | 0.042 |
| 2.000 | 0.166 | 0.313 | 5.776 | 0.016 |
| 2.246 | 0.166 | 0.231 | 2.225 | 0.136 |
| 2.838 | 0.166 | 0.212 | 1.881 | 0.170 |
| 3.000 | 0.166 | 0.227 | 1.448 | 0.229 |

| Distance | PVEself | PVEneighbor | $\chi_1^2$ | p-value |
| --- | --- | --- | --- | --- |
| 3.172 | 0.166 | 0.144 | 0.551 | 0.458 |
| 3.616 | 0.166 | 0.137 | 0.524 | 0.469 |
| 4.000 | 0.166 | 0.093 | 0.150 | 0.699 |
| 4.133 | 0.166 | 0.079 | 0.091 | 0.762 |
| 4.253 | 0.166 | 0.060 | 0.044 | 0.833 |

(C) Scald
| Distance | PVEself | PVEneighbor | $\chi_1^2$ | p-value |
| --- | --- | --- | --- | --- |
| 0.000 | 0.511 | 0.000 | 14.292 | 1.6E-04 |
| 1.000 | 0.511 | 0.224 | 27.382 | 1.7E-07 |
| 1.424 | 0.511 | 0.118 | 13.395 | 2.5E-04 |
| 2.000 | 0.511 | 0.200 | 15.411 | 8.6E-05 |
| 2.246 | 0.511 | 0.128 | 7.043 | 0.008 |
| 2.838 | 0.511 | 0.124 | 5.958 | 0.015 |
| 3.000 | 0.511 | 0.158 | 8.134 | 0.004 |
| 3.172 | 0.511 | 0.159 | 6.784 | 0.009 |
| 3.616 | 0.511 | 0.171 | 7.391 | 0.007 |
| 4.000 | 0.511 | 0.193 | 9.314 | 0.002 |
| 4.133 | 0.511 | 0.202 | 9.599 | 0.002 |
| 4.253 | 0.511 | 0.193 | 7.883 | 0.005 |

